# Conjunctive representation of what and when in monkey hippocampus and lateral prefrontal cortex during an associative memory task

**DOI:** 10.1101/709659

**Authors:** Nathanael A. Cruzado, Zoran Tiganj, Scott L. Brincat, Earl K. Miller, Marc W. Howard

## Abstract

Adaptive memory requires the organism to form associations that bridge between events separated in time. Many studies show interactions between hippocampus (HPC) and prefrontal cortex (PFC) during formation of such associations. We analyze neural recording from monkey HPC and PFC during a memory task that requires the monkey to associate stimuli separated by about a second in time. After the first stimulus was presented, large numbers of units in both HPC and PFC fired in sequence. Many units fired only when a particular stimulus was presented at a particular time in the past. These results indicate that both HPC and PFC maintain a temporal record of events that could be used to form associations across time. This temporal record of the past is a key component of the temporal coding hypothesis, a hypothesis in psychology that memory not only encodes what happened, but when it happened.

## Introduction

Many studies (Jones & Wilson, 2005; Hyman, Zilli, Paley, & Hasselmo, 2005, 2010; H. Kim, 2011) show that interactions between hippocampus (HPC) and prefrontal cortex (PFC) are essential to forming associations between unrelated stimuli. While early conceptions of associative memory treated association as a scalar atomic value (Watson, 1913; Clark, Lansford, & Dallenbach, 1960; Hawkins & Kandel, 1984), other models of memory incorporated a temporal record of the past as a key component in learning associations— learning not only the association between the stimuli but also the order and timing (James, 1890; Arcediano & Miller, 2002).

Stimulus selective time cells could serve as the neural mechanism by which a temporal record is maintained and updated. Time cells are neurons that fire sequentially, each for a circumscribed period of time, during the delay interval of a memory task (Pastalkova, Itskov, Amarasingham, & Buzsaki, 2008; MacDonald, Lepage, Eden, & Eichenbaum, 2011), provide a neural representation that includes information about time. Time cells have been observed extensively in mice and rats (Howard et al., 2014; Salz et al., 2016) and they have also been observed in primate PFC (Tiganj, Cromer, Roy, Miller, & Howard, 2018; Jin, Fujii, & Graybiel, 2009). Recent studies have shown temporal coding in primate HPC (Sakon, Naya, Wirth, & Suzuki, 2014; Naya & Suzuki, 2011; Naya, Chen, Yang, & Suzuki, 2017) suggesting that time cells may be found in this region as well. Although these results are suggestive, it has not been definitively shown if time cells exist in monkey HPC and if these time cells are similar in their proprieties to monkey PFC time cells.

We analyze neural recording from monkey HPC and PFC during a memory task that requires the monkey to associate stimuli separated by about a second. After the first stimulus was presented, a large number of units in both HPC and PFC fired in sequence. Many units fired only when a particular stimulus was presented at a particular time in the past. These results indicate that both HPC and PFC maintain a temporal record of the past. This temporal record of the past is consistent with the idea that time is intrinsic to forming associations Arcediano and Miller (2002).

## Methods

This paper reports new analyses of data previously published in Brincat and Miller (2015). In Brincat and Miller (2015), monkeys performed a visual paired associate task while recordings were taken of neural activity in HPC and PFC. This paper introduces single unit analyses to classify and characterize neurons with temporal firing fields and a population level analysis of temporal information intended to support the single unit analyses.

### Task

Two macaque monkeys were trained to perform a visual paired associate task. The monkeys were trained on the task for several weeks prior to recording. For each recording session, six visual stimuli were chosen at random, with four serving as cue objects (labeled as A, B, C and D for the purposes of this paper) and two as associate objects (labeled X and Y). Each trial started with fixation on a white dot for 500 ms, followed by cue presentation for 500 ms, followed by a delay for 750 ms, followed by either a match or non-match stimulus for 500 ms, followed by reward for a correct response to match stimulus, no reward for incorrect responses, and an additional delay of 500 ms followed by match stimulus for correct responses to non-match stimulus (Figure 1). Trials without a valid response (i.e. the monkey did not maintain fixation) were discarded from the data set. Each session consisted of 36 training trials where the stimuli were passively presented followed by 96 identity match to sample trials in which the same stimulus was presented for cue and match in order to familiarize the monkey with the stimuli, followed by the actual test trials (typically around 1000 trials for each session). The analyses in this paper only use the data from the test trials, not the training trials or identity match to sample trials.

**Figure 1.**
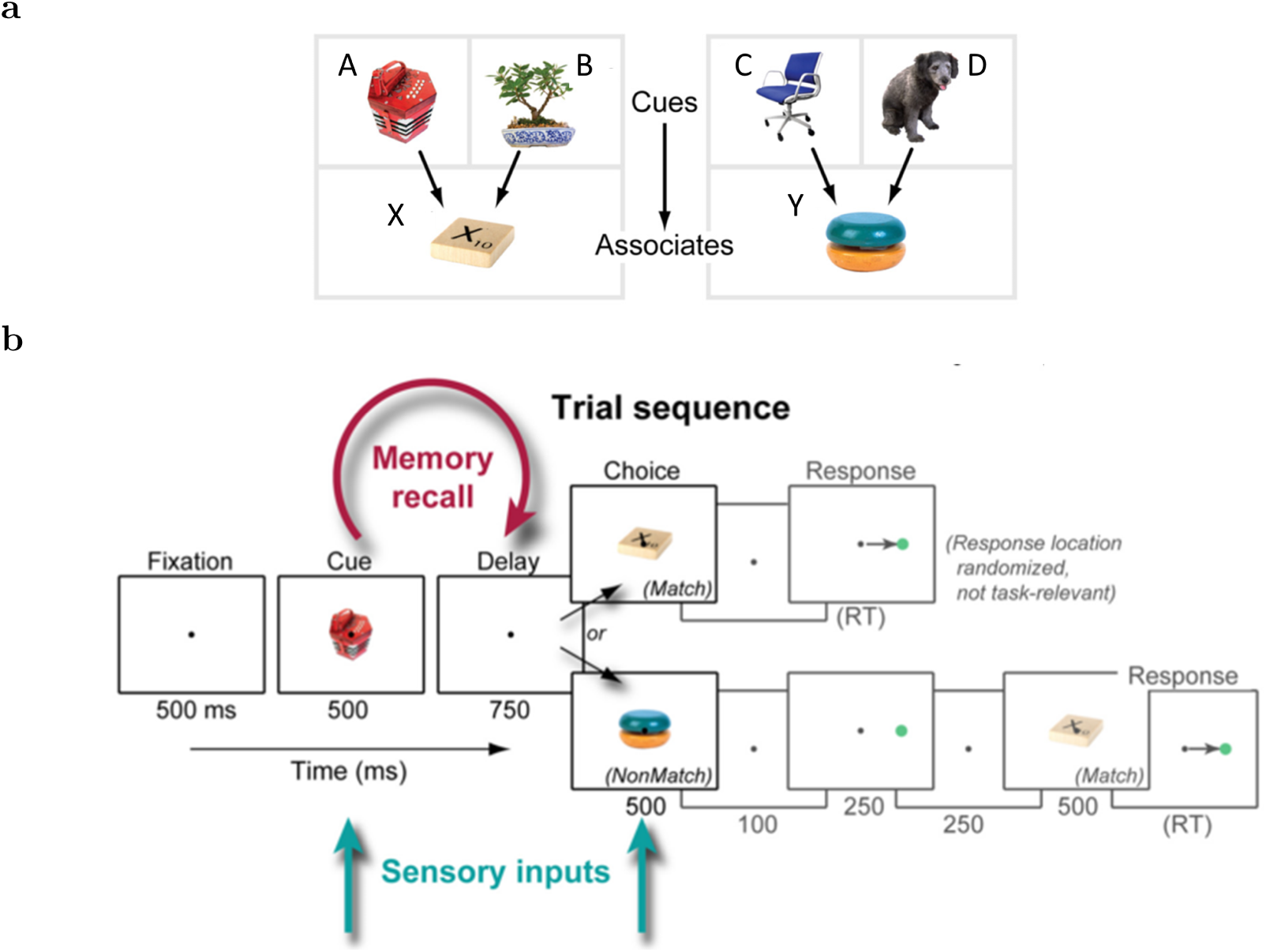
The experimental design required the monkey to use associative memory. The correct response depended on both learning an association and retaining a stimulus identity in working memory. **a:** Monkeys were trained in a paired associate task. For each recording session the monkey learned two pairs of cue stimuli (A/B and C/D) and a corresponding associate stimulus for each pair (X and Y). **b:** Sequence of events in a trial. The identity of the cue stimulus determines the identity of the match stimulus. The analyses in this paper focus on the 1250 ms from initial cue presentation to end of the delay period.

For the purposes of identifying time cells we analyze the 1250 ms from cue presentation until match or non-match stimuli presentation. We restrict our analyses to the cells that were identified as properly isolated, as classified by Brincat and Miller (2015).

### Recording

The recording techniques are described in detail in Brincat and Miller (2015). To summarize, multiple microelectrodes were lowered daily into prefrontal cortex and hippocampus. Specifically, recording sites included all subregions of the hippocampus (CA1, CA2, CA3, dentate gyrus, subiculum) and dorsolateral and ventrolateral PFC (including parts of areas 46, 45, and 8). The microelectrodes were referenced to ground.

For each session, up to 16 electrodes were inserted into the PFC and up to 4 electrodes were inserted into the HPC. The spikes were recorded using epoxy-coated tungsten electrodes (FHC) amplified by a high-input-impedance headstage filtered to extract spiking activity, and threshold-triggered to separate neuronal spikes from background noise. The spiking signal was not prescreened for task responsiveness or selectivity in order to avoid introducing bias. The electrodes were targeted with custom Matlab software.

#### Data Selection

For each testing trial we analyzed the 1250 ms starting from presentation of the cue stimulus and terminating at the presentation of the associate stimulus. This interval includes the 500 ms presentation of the cue stimulus and a 750 ms blank delay interval. Only data from the testing trials were used, none of the training trials or identity match trials were used in this analysis. As a preprocessing step to both the maximum likelihood analysis and the linear discriminant analysis, spike trains were downsampled to 1 ms temporal resolution such that if a spike was observed in a particular 1 ms time bin, the corresponding data point were set to 1, otherwise it was set to 0.

### Single Unit Analysis

The goal of the analysis used in this paper is to classify and characterize neurons with temporal firing fields. We classified these neurons as time cells. This analysis is designed to be robust to single trial variability and minimize the number of arbitrary parameters used, while still providing insight into individual cells.

#### Maximum Likelihood Analysis

These methods build on analysis methods used to classify time cells in rodent hippocampus and mPFC (Salz et al., 2016; Tiganj, Kim, Jung, & Howard, 2017) and stimulus selective time cells in monkey lPFC (Tiganj et al., 2018).

As in (Tiganj et al., 2018), this analysis optimized parameters of a Gaussian receptive field to best fit the firing rate of each potential time cell. We used maximum likelihood parameter estimation using a combination of gradient descent optimization and particle swarm. Nested statistical models can be compared with a likelihood ratio test. The likelihood ratio test (Wu, 1999) incorporates the difference in the number of parameters of the nested models as the degrees of freedom in a χ^2^ test. We first trained the parameter of a constant firing rate model as a control, then a Gaussian receptive field (Gaussian time field plus constant rate), then additional models with parameters for stimulus selectivity.

#### Fitting the Parameters

The spike trains of each cell were fitted with different models that allowed the firing rate to vary with different variables, such as time and stimulus identity. The parameter space of these models was systematically explored in order to compute the maximum likelihood fit. To find the best-fitting model the parameter space was iteratively searched using a combination of particle swarming and the QuasiNewton method. Particle swarming was performed first (with the swarm size equal to 50) and its output was used to initialize the Quasi-Newton method which was performed second (the number of maximum function evaluations was set to 10000). The computations were implemented in Matlab 2016a. To avoid solutions that converged to a local minimum, the fitting procedure was repeated until the algorithm did not result with better likelihood for at least ten consecutive runs.

The likelihood of the fit was defined as a product of these probabilities across all 1250 time bins within each trial and across all trials. We expressed the likelihood in terms of the negative log-likelihood (nLL), therefore instead of a product, a sum of the terms was computed:

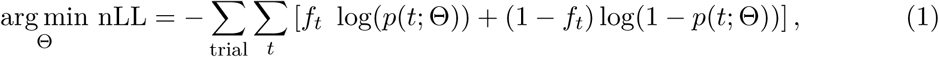

where *f*_*t*_ is the spike train, Θ is the parameters of the model.

#### Constant Firing Rate

We first fitted a constant firing rate model to serve as a control. The likelihood was set to the constant term *a*_0_.

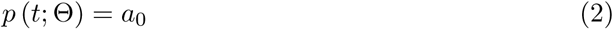

#### Gaussian Receptive Field

To estimate temporal variability in firing, we augmented the constant firing rate model (eq. 2 with a term describing a Gaussian time field:

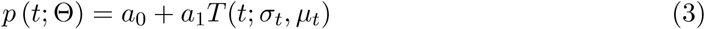

where the Gaussian-shaped time field *T* (*t*; *σ*_*t*_, *µ*_*t*_) was defined as:

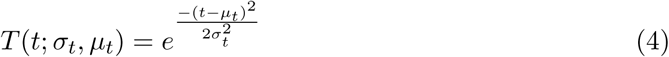

The Gaussian receptive field described by Equations 3 has an additional three parameters compared to the constant firing rate model. In addition to the background firing rate *a*_0_, this model includes parameters for the amplitude of the time field *a*_1_, and the mean *µ*_*t*_ and standard deviation *σ*_*t*_ of the time field.

The mean of the time term *µ*_*t*_ was allowed to vary between -4375 ms and 5625 ms and the standard deviation *σ*_*t*_ varied between 10 and 10000 ms. This range of parameters can not only account for time cells, but also describe cells with monotonically changing firing rates. When *µ*_*t*_ is less than 0 ms, the Gaussian receptive field describes a cell with monotonically decreasing firing rate. Likewise, when *µ*_*t*_ is greater than 1250 ms the Gaussian receptive field describes a monotonically increasing firing rate. In order to ensure that *p* (*t*; Θ) can be considered as a probability we had to ensure that its values are bounded between 0 and 1. Therefore, the coefficients were bounded such that *a*_0_ + *a*_1_ ≤ 1. *a*_*n*_ was not allowed to go below 0.

#### Stimulus Specific Gaussian Receptive Field

The next statistical model included additional parameters to allow the time-varying component to depend on the identity of the stimulus that began the trial. Stimulus specificity was tested with a model that allowed four parameters, rather than one as above, to modulate the Gaussian-shaped time field. The probability of a spike at time point *t* was given as:

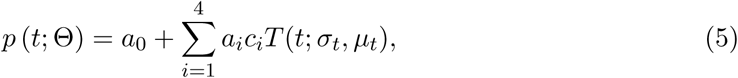

where *a*_0_ to *a*_4_, *µ*_*t*_, and *σ*_*t*_ are the parameters to be estimated. The factor *c*_*i*_ was equal to 1 for trials when a stimulus from *i*-th cue stimulus was presented and 0 otherwise. For instance, *c*_1_ = 1 for trials which started with stimulus A and *c*_1_ = 0 for trials where the sample stimulus was cue stimuli B, C or D).

#### Log Likelihood Test

In order to quantify whether the contribution of the terms that contained time was significant, the maximum log-likelihoods were compared. Note that the constant term model is equivalent to the Gaussian receptive field modulated to zero (*a*_1_ = 0). Since the models with and without the Gaussian receptive field are nested, the likelihood-ratio test could be used to assess the probability that adding the time term significantly improved the fit. The test is based on the ratio of the likelihoods of two different statistical models and expresses how many times the data are more likely under one model than the other and it takes into account the difference in the number of parameters. We refer to cells that were better fit by Eq. 3 than by a constant term (just *a*_0_) as *time cells*, subject to several additional constraints. To ensure that a unit will not be classified as a time cell only due to its activity in a single trial and to correct for other possible trial averaging errors, the analysis was done separately on even and odd trials. For a unit to be classified as a time cell it was required that the likelihood-ratio test was significant (*p* < 0.01) for both even and odd trials. In order to eliminate units with monotonically changing firing rate during a delay interval, *µ*_*t*_ was required to be within the delay interval and at least one *σ*_*t*_ away from either the beginning or the end of the interval for time cells. Units outside these boundaries were either monotonically changing (ramping/decaying) cells or ambiguous cells. Units with *µ*_*t*_ entirely outside the 0-1250 ms interval were classified as monotonic changing cells, and units within 0-1250 ms but within one *σ*_*t*_ of these bounds were classified as ambiguous cell.

The model that includes stimulus specificity eq. 5 and the model with a single time field eq. 3 are nested. Therefore, we use the likelihood-ratio test to assess the probability that adding the stimulus specificity significantly improves the fit. When the outcome of the likelihood ratio test was significant (*p* < 0.01), a time cell was classified as stimulus specific.

### Linear Discriminant Analysis

#### Overview

To quantify decoding accuracy at a population level we used linear discriminant analysis (LDA). We divided the 1250 millisecond interval composed of presentation and delay period into 50 ms long non-overlapping time bins. For each region we used the entire population of neurons. The units were recorded during multiple recording sessions with different number of trials. To ensure an equal number of trials across all units we restricted the number of trials to the lowest number of trials recorded from a single unit (606 trials). For each time bin we trained an LDA classifier on 80% of randomly chosen trials and used the remaining 20% of the trials for testing. The LDA classifier was implemented using Matlab 2016a function *classify*. To ensure stability of LDA the dimensionality of the training data was reduced to full rank by using the Matlab function rref to identify the pivots before each run of the classifier. The matrices need to be full rank in order to run the classification procedure because the LDA needs to use the matrix inverse.

#### Population Stimulus Specificity

To assess stimulus specificity at the population level, the classifier was trained to assign each trial to one of the four stimuli (classifying a trial as A, B, C, or D). The testing classification was done on all time bins (to evaluate performance of the classifier as a function of temporal distance between training and testing time bin). We repeated the training and testing for 10 iterations (taking random subsamples of the 606 trials) in order to obtain robust results (quantified through standard error of the mean).

## Results

To anticipate the results, the maximum likelihood analysis identified many units as time cells and stimulus specific time cells in both HPC and PFC. Fewer units were identified as ambiguous cells (possibly ramping cells or possibly time cells) and a small number of units were identified as monotonically changing cells. HPC and PFC time cells were qualitatively similar for several properties, suggesting that HPC and PFC share a common temporal code. Both populations of time cells were temporally compressed. Evidence of stimulus selectivity, temporal compression, and encoding the past, not the future, also existed at the population coding level, as confirmed with linear discriminant analysis.

### Conjunctive coding of time and stimulus identity

A likelihood ratio test was used to compare models for the firing rate of each unit. Units which were significantly better described by the time cell model than by the constant firing rate model are referred to as time cells. Units that were significantly better described by the stimulus specific time cell model than by the time cell model are referred to as stimulus specific time cells. Examples of units identified as stimulus specific time cells are shown in Figure 2 for HPC and Figure 3 for PFC. These units have a variety of firing field peak times, firing field widths, and overall firing rate and respond to all combinations of stimuli.

**Figure 2.**
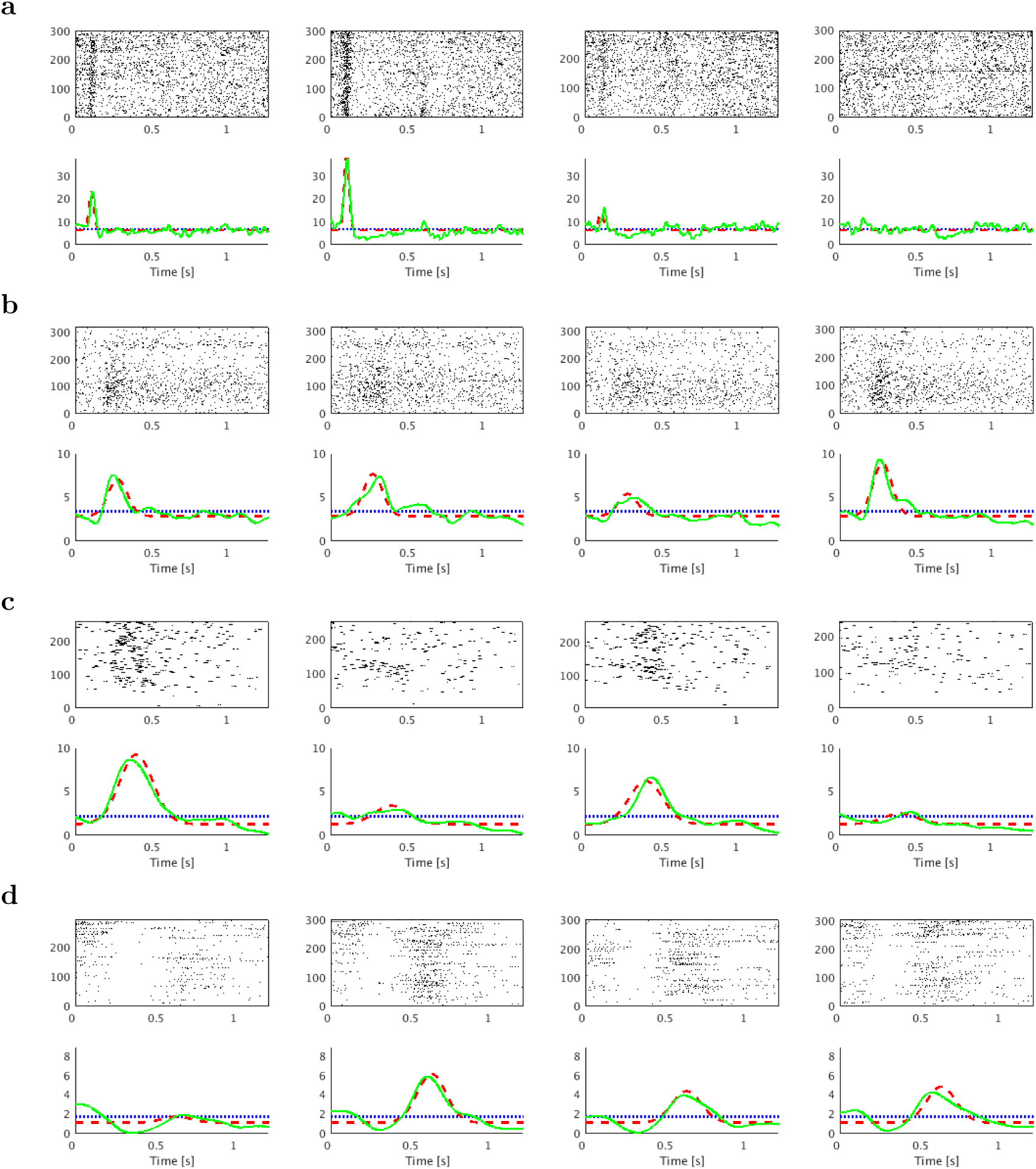
Examples of HPC cells that were conjunctively selective for stimulus identity and time. Each of the four columns correspond to trials with a different cue stimulus and **a-d** are different hippocampal cells. Within each raster each row is a trial, and time is shown from cue stimulus onset to associate stimulus onset. Within each graph, the green line is the smoothed firing rate, the blue line is a fitted constant firing rate, and the red line is the fitted stimulus specific time cell model. Potential time cells were characterized by increased firing rate at a particular time. The time cells in **a** and **b** responded earlier in the delay interval, the time cells in **c** and **d** responded later in the delay interval. The unit in **a** is selective for stimuli A and B. The unit in **b** is selective for stimuli A, B, and C. The unit in **c** is selective for stimuli A and C. The unit in **d** is selective for stimulus B.

**Figure 3.**
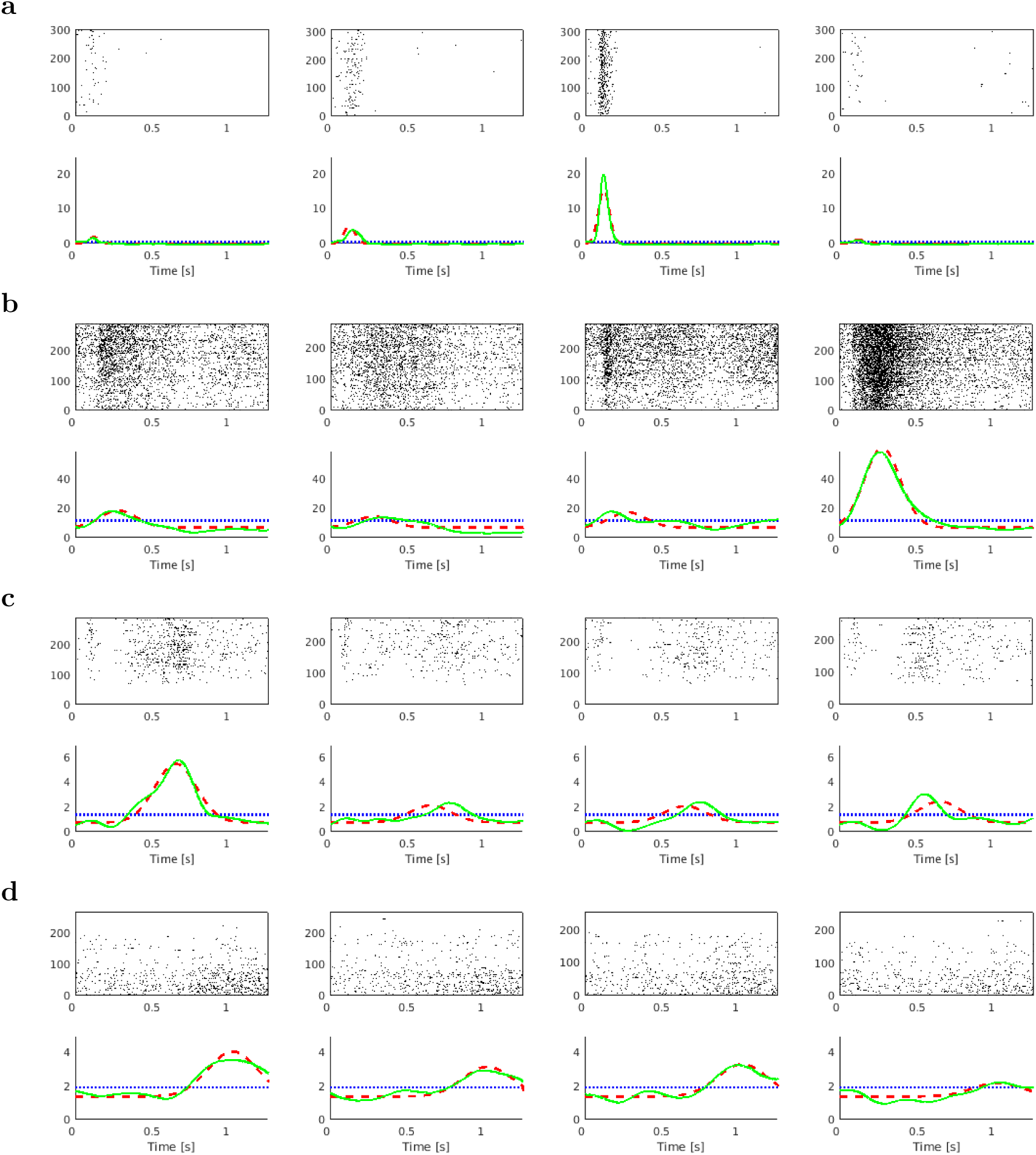
Examples of PFC cells that conjunctively coded for stimulus identity and time. As in Figure 2, each of the four columns correspond to trials with a different cue stimulus and **a-d** are different cells. Within each graph, the green line is the smoothed firing rate, the blue line is a fitted constant firing rate, and the red line is the fitted stimulus specific time cell model. Potential time cells were characterized by increased firing rate at a particular time. The time cells in **a** and **b** responded earlier in the delay interval, the time cell in **c** towards the middle of the interval and the time cell in **d** responded later in the delay interval. The unit in **a** is selective for stimuli C. The unit in **b** is selective for stimulus D. The unit in **c** is selective for stimulus A. The unit in **d** is selective for stimuli A, B, and C.

Overall a substantial number of units were identified by the maximum likelihood method: 246 units for PFC and 125 units for HPC clearly met our criteria as time cells and 149 units for PFC and 68 units for HPC met our criteria as stimulus specific time cells. Thus, both HPC and PFC support conjunctive representations of time and stimulus identity. Although the proportion of time cells in HPC (.29) vs PFC (.43) is reliably different (the proportions differ with significance *p* < .001 according to a two proportion z-test), this difference is not necessarily meaningful, but could be a confound of the overall difference in firing rates between the two areas. All else equal, more spikes leads to improved resolution of a putative time field. All else equal, higher firing rates do not lead to more false positives, but lower firing rates do lead to more false negatives. Comparing HPC and PFC firing rate average across the 1250 millisecond delay interval for each cell using a Wilcoxon rank sum test shows that HPC cell median firing rate is significantly less than PFC median firing rate (HPC median: .22 Hz HPC mean: 1.40 Hz, PFC median: .44 Hz, PFC mean: 1.73 Hz, z-statistic -4.02, rank sum 2.33*e*5, p<.001). This higher firing rate is consistent with the fact that overall a higher proportion of PFC cells were identified as time cells.

### Time cells were the primary form of temporally modulated cells

As an additional control on the time cell analysis similar methods were used to look for cells with ramping and decaying firing rates, i.e. cells with monotonically changing firing rates. Compared to the units clearly classified as time cells, a fewer number of units were ambiguously between proper time cells and monotonically changing cells, 74 units for PFC 34 units for HPC. The units were not included in the analysis of time cells shown in Figure 4, but are included in Supplemental Figure S1. Even with the ambiguous cells included, the overall results showing temporal compression are still present.

**Figure 4.**
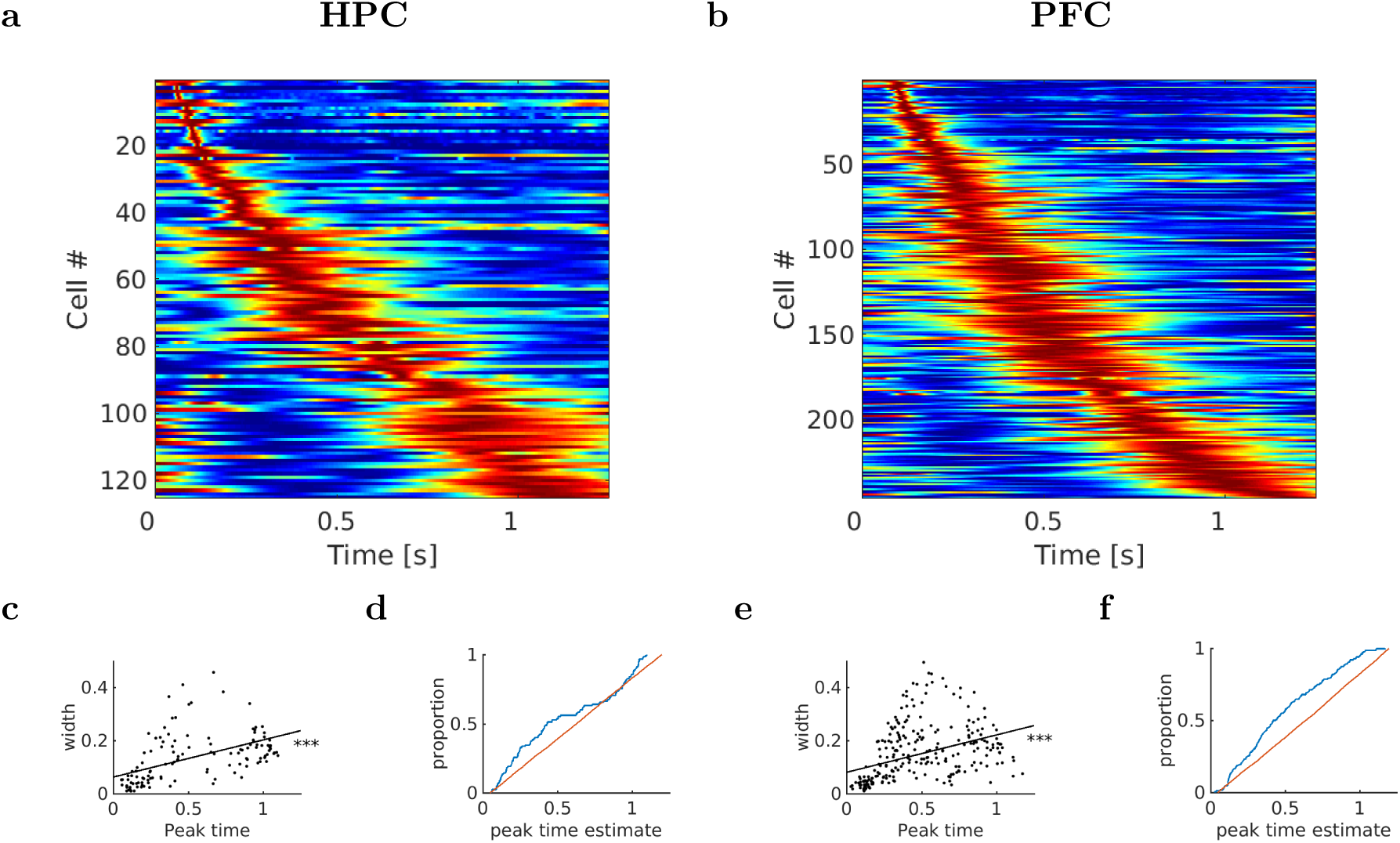
Time cells in HPC (left) and PFC (right) tiled the interval in a compressed timeline. **a-b:** Activity of 125 HPC units and 246 PFC units classified as time cells. Each row corresponds to a single unit and displays the firing rate (normalized to 1) averaged across all trials. Red corresponds to high firing rate; blue corresponds to low firing rate. The time cells are sorted with respect to the peak time estimated for their time field. There are two features related to temporal accuracy that can be seen from examination of these figures. First, time fields later in the delay were more broad than time fields earlier in the delay (see also **c** and **e**). This can be seen as the widening of the central ridge as the peak moves to the right. In addition the peak times of the time cells were not evenly distributed across the delay, with later time periods represented by fewer cells than early time periods (see also **d** and **f**). This can be seen in the curvature of the central ridge; a uniform distribution of time fields would manifest as a straight line. **c:** Width of the time fields as a function of the peak time in HPC. Each dot represents the best-fitting parameters for a single unit classified as a time cell. The blue line is a fitted linear model. The linear regression is significant *p* < .001. **d:** Peak times of the time fields in HPC are non-uniformly distributed along the delay interval. The blue line is the cumulative distribution function of the time cell peak times. The red line is a uniform distribution. The distributions differ at *p* < .001 as estimated from a Kolmogorov-Smirnov test. **e:** Same as **c** but for time cells in PFC. The linear regression is significant at *p* < .001. **f:** Same as **d** but for time cells in PFC. Kolmogorov-Smirnov test is significant at *p* < .001.

Only a small number of units were classified as monotonically cells (14 for HPC and 22 for PFC). This is significantly less than the number of time cells (significant at p<.01 by a two proportion z-test). Additionally, this analysis doesn’t rule out other possible forms of firing rate modulation such as a time cell with a peak outside the allowed interval. Example monotonic cells are shown in the Supplemental, (Figures S2 and S3). The monotonic cells were not as consistent or reliable, even the best examples of monotonic cells show irregularities. Overall, monotonically changing cells were substantially less common than time cells.

### Temporal Compression

The Figure 4 heatplots show several key patterns in the HPC and PFC time cells. The width of the central ridge in the heatplots increases from the left of the plot to the right of the plot, suggesting that time cells that fire earlier in the trial tend to have narrower time fields than the time cells that fire later. The central ridges in the heatplots do not follow a straight line, as would have been the case if it followed a uniform distribution. Rather, the curve flattens as the interval proceeds. Qualitatively, HPC and PFC were similar in that they both show the overall characteristics of a compressed representation of time.

A compressed representation refers to a neural representation that does not have the same resolution across the entire range it is representing. A compressed representation of time does not have constant resolution across the range of time it represents. Time cells with a constant resolution would have uniform firing field widths with uniformly spread peak times across the elapse of time during the trial. If the neural representation of time in this data set was not constant, there are two ways the identified time cell population could show temporal compression. First, the width of time fields would increase as the trial elapses. Second, the number of time cells with time fields earlier in the delay would be larger than the number later in the delay. Both of these properties were qualitatively observed in the heatplot, as discussed above, and were confirmed with quantiative measures, discussed below. To provide an intuition into the meaning of these measures, Supplemental Figure S5 shows the results of applying these analyses to simulations of ideal time cells.

#### Time cell receptive field width increases with peak time

The first qualitative impression of the heatplot can be confirmed by analyses of the across-unit relationship between the peak time (*µ*_*t*_) and the standard deviation (*σ*_*t*_) of the estimated Gaussian shaped time fields (Figure 4). This across-unit relationship has a positive, statistically significant slope for both HPC (intercept .06 ±.009 s *p* < .001, slope .14 ±.015 s/s *p* < .001, *r*^2^ = .24) and PFC (intercept .08 ±.007 s *p* < .001, slope .14 ±.013 s/s *p* < .001, *r*^2^ = .16). Including ambiguous cells, the statistical significance is much weaker, but there is still a trend in HPC (intercept .15 ±.026 s *p* < .001, slope .07 ±.039 s/s *p* < .1) and PFC (intercept .06 ±.017 s *p* < .001, slope .06 ±.027 s/s *p* < .05, *r*^2^ = .16). Time cells firing earlier have narrower time fields in both HPC and PFC. These findings are consistent with a key quantitative prediction of a compressed timeline.

#### Distribution of time cell receptive fields show temporal compression

The cumulative density functions of time cell centers in HPC and PFC are shown in Figures 4d and 4f respectively. We compared the empirical distribution of the peak times of the time cells to a uniform distribution using a Komolgorov-Smirnov test. The KS test rejected the hypothesis that the distribution of the peak times is uniform for both HPC (using only time cells *n* = 125, ks-stat=0.23, *p* < .001; using time cells and ambiguous cells *n* = 159, ks-stat=0.18, *p* < .001) and PFC (using only time cells *n* = 246, ks-stat=0.02, *p* < .1; using time cells and ambiguous cells *n* = 320, ks-stat=0.09, *p* < .001). These results demonstrate that more time cells code for points earlier in the interval than later in the interval in both HPC and PFC. This implies decreasing temporal accuracy as the delay proceeds.

PFC and HPC time cell peak times distributions differ from each other as shown by a KS test (KS stat .1151, p<.05). HPC time cells tended to peak earlier in the interval in comparison to PFC time cells. Although this difference is statistically significant, in absolute terms it is small. The cause and meaning of this difference is unclear. One possible explanation is that although the two regions carry similar temporal signal (as evidenced by the overall similar temporal compression), the latency for this signal to manifest is smaller in HPC than PFC.

### Population Analysis

Linear discriminant analysis (LDA) was used to investigate the temporal dynamics of the stimulus identity coding on the population level. LDA was performed to decode stimulus identity of the cue stimulus at each 50 ms time bin of the sample and delay intervals (see methods). LDA was able to decode stimulus identity above chance (at p<.01 significance, as measured by a one proportion z-test on each bin separately) for HPC and PFC. Additionally, the decoding accuracy along the diagonal during the 1250-ms period was higher (at p<.01 significance, as measured by a two proportion z-test on each bin separately) than the entire rest of the 1250-ms period for both HPC and PFC (shown with the contour lines in Figure 5). If stimulus identity were represented via persistent firing cells that did not change their firing rate during the delay, the decoder would have similar accuracy over the entire period of time following onset of the cue. This indicates that temporal representation is conjunctive with stimulus identity. The decoding accuracy is also higher towards the beginning of the 1250 ms period (at p<.01, as measured by a two proportion z-test on each bin separately), showing temporal compression (again, shown with the contour lines in Figure 5). These results are consistent with the results of the single unit analyses.

**Figure 5.**
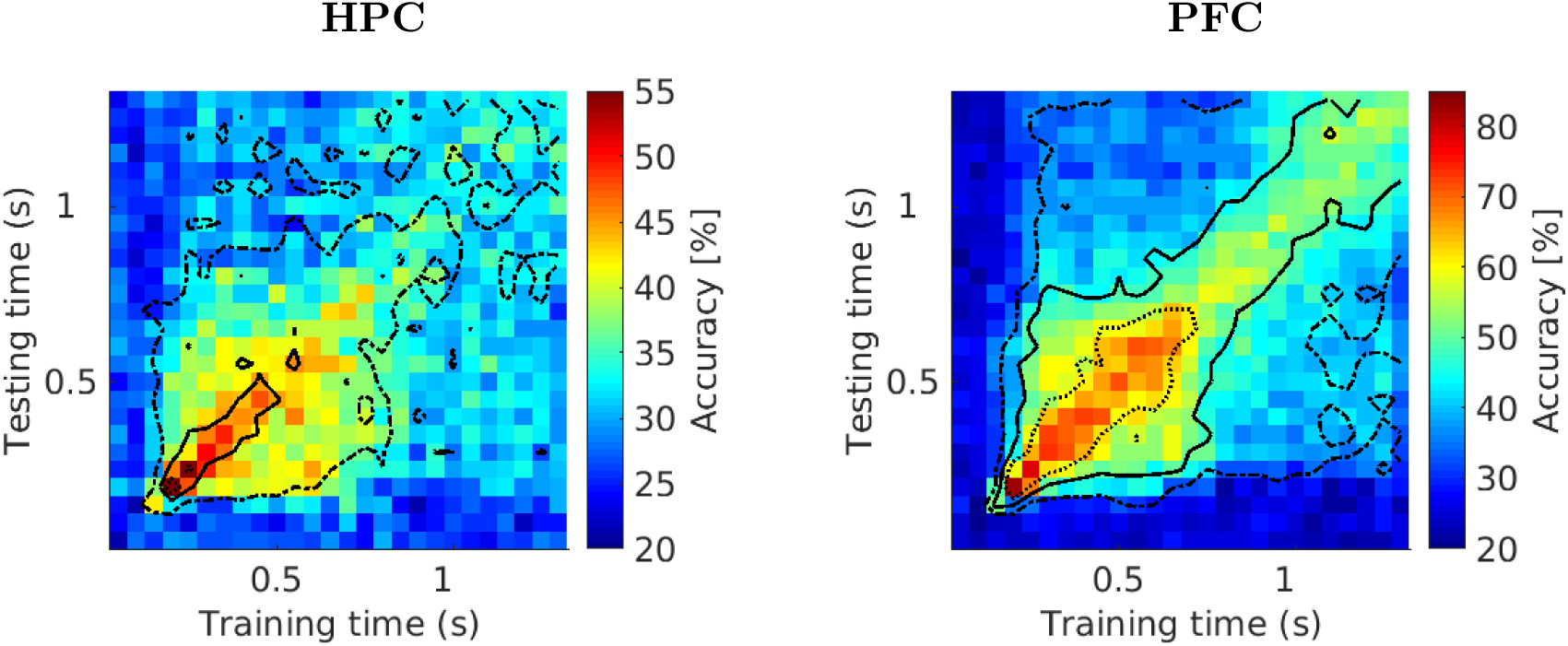
Ensembles in HPC and PFC can decode what happened when. displays the accuracy of a linear discriminant analysis (LDA) cross-temporal classifier applied on 50 ms time bins. Each bin provides classification accuracy for the classifier trained on a time bin denoted on x-axis and tested on a time bin denoted on y-axis. 80% of trials were used for training with 20% of trials held out for testing. Note that the colorscale is different between panels. Overall accuracy was higher for PFC than HPC, but there are also a higher number of cells avaiable in PFC. Each contour represents an additional level of significance compared to the previous contour (at p<.01). Together, these contours quantitatively illustrate that the stimulus identity is encoded, that this encoding is temporally dependent, and that this encoding is a compressed representation.

## Discussion

Time cells are neurons that fire for a circumscribed period of time after a defined event. A population of time cells collectively spans the interval with activity. Time cells might serve an important role in maintaining a temporal record of the past. Previous research has identified time cells in rodents (e.g., Pastalkova et al., 2008; MacDonald et al., 2011). Naya and Suzuki (2011) found temporally modulated firing rates with stimulus selectivity in primate PFC, but did not specifically look for Gaussian receptive fields. In this study, we found primate HPC and PFC both had a substantial number of time cells with several key properties. The time cells in both of these areas were stimulus specific, allowing decoding of the stimulus presented at the beginning of the delay interval. The time cells in both HPC and PFC showed the collective proprieties of a compressed representation (Figure 4). Population analyses yielded similar conclusions (Figure 5). Collectively, these results show that HPC and PFC encode a compressed temporal record of what happened when. This record decreased in temporal resolution as the stimulus receded into the past.

### A common temporal code for associative memory

The analyses presented here have showed that HPC and PFC share a common temporal code with both stimulus selectivity and temporal compression. The spiking activity contained information of not only when stimuli happened, but what stimuli happened. This record of what happened when is a key component of the temporal coding hypothesis, the idea that associative memory includes not only pairs of associated stimuli but also their temporal relationships (Arcediano & Miller, 2002). This record of what happened when also showed temporal compression. This temporal compression was characterized by two key proprieties. First, time fields later in the sequence were more broad (i.e., less precise). Second, there were more neurons with time fields early in the delay and fewer neurons representing times further in the past.

The stimulus selectivity found in the time cells identified here is consistent with broader hypotheses about mixed selectivity. The term mixed selectivity is used to describe responses across a population of neurons to diverse, non-linear combinations of task relevant variables. Mixed selectivity results in a high dimensional neural representations which enables simple readouts to generate a high number of responses (Fusi, Miller, & Rigotti, 2016; Rigotti et al., 2013). Mixed selectivity has been found in a variety of time cells populations for a variety of task relevant variables, including place (MacDonald et al., 2011; Tiganj, Cromer, Roy, Miller, & Howard, 2017; Salz et al., 2016), cue stimulus identity (Terada, Sakurai, Nakahara, & Fujisawa, 2017; Tiganj et al., 2018), and motor actions (Jin et al., 2009; Mello, Soares, & Paton, 2015). The results of this paper add to these previous results by showing mixed selectivity in response to a visual task paired associate task in primate HPC and PFC simultaneously.

The temporal compression identified in this report is consistent with past work, both behavioral work and previous reports of time cells. Behavioral work on timing shows that the accuracy in estimating the elapsed time decreases with the amount of time to be estimated (Rakitin et al., 1998; Lewis & Miall, 2009). Temporal compression is consistent with this decrease in temporal accuracy. Temporal compression has been observed, for time cells in a variety of brain regions, including the hippocampus (Howard et al., 2014; Salz et al., 2016), entorhinal cortex (Kraus et al., 2015), medial prefrontal cortex (mPFC) (Tiganj, Kim, et al., 2017), lateral prefrontal cortex (lPFC) (Tiganj et al., 2018), dorsolateral prefrontal cortex (dlPFC) (Jin et al., 2009), and striatum (Jin et al., 2009; Mello et al., 2015; Akhlaghpour et al., 2016). The results of this paper add to past results by showing cue stimuli selectivity and temporal compression in primate HPC simultaneously with PFC.

This observed temporal compression is consistent with the classical behavioral observation that learning is invariant of a temporal scale. Scale invariance is the property that the features of a system do not change so long as the scales involved are multiplied by a common factor. For instance, Balsam and Gallistel (2009) reviewed evidence that the learning of an association between a conditioned stimulus and unconditioned stimulus was not dependent on the time between them, but rather the ratio of the time between them and the overall time from trial to trial. Thus when both delay and intertrial interval were rescaled by a same factor, the number of trials that the animal needed to learn the pairing between conditioned and unconditioned stimulus remained constant, indicating that the leaning process is invariant of the time scale. In other words, the process of associative learning was not dependent on some critical window of time, but rather can work across a variety of scales so long as the relevant timings are proportionate. A compressed temporal code could potentially explain how this scale invariance is constructed, if the resolution of the temporal code scales proportionality across the time interval it encodes, this representation would be scale invariant (Tiganj, Cromer, et al., 2017; Shankar & Howard, 2013). The results reported here showed a temporal code with non-uniform resolution (higher accuracy in the beginning of the interval), allowing for the possibility that the resolution of the time code might scale proportionality to the elapsed time in primate HPC and PFC.

Previous studies have described temporal coding schemes with increasing firing rates as a duration elapses (J. Kim, Ghim, Lee, & Jung, 2013; Sakon et al., 2014) and increasing dopamine concentration as a reward approaches (Howe, Tierney, Sandberg, Phillips, & Graybiel, 2013). Note that ramping firing rates are not mutually exclusive with time cells. A time cell with a peak at the time of a reward would have the same pattern of firing rate as ramping cell (over the delay period between cue presentation and reward). For the purposes of this study, however, we have defined time cells to exclude this possibility to get a conservative estimate of the number of time cells. Similarly, there have been reports of firing rates that decay as a duration elapses (J. Kim et al., 2013; Sakon et al., 2014; Blanchard, Strait, & Hayden, 2015). In particular, decaying and ramping of an exponential form is described in Monkey EC in Bright et al. (2019). Notably, a Gaussian receptive field is nested with the statistical model used by Bright et al. (2019) and yet they did not identify any cells best described by a Gaussian receptive field, i.e. they did not identify any time cells. It is possible that differences in experimental paradigms can result in different temporal coding schemes taking precedent.

### Persistent Firing and Dynamic Coding for Working Memory

We found a large number of cells that coded for the identity of past stimulus but only fired for a short period of time. Nonetheless, the population of such cells enabled readout of stimulus identity throughout the delay. This pattern is not predicted by classical models about the maintenance of information in a fixed-capacity buffer (Atkinson & Shiffrin, 1968) that lead to the idea that the brain maintains information *via* persistent stimulus-specific firing that is not temporally modulated (Fuster & Alexander, 1971; Funahashi, Bruce, & Goldman-Rakic, 1989; Goldman-Rakic, 1996). This view led to computational models that predict persistent neural activity (e.g., Compte, Brunel, Goldman-Rakic, & Wang, 2000; Durstewitz, Vittoz, Floresco, & Seamans, 2010; Chaudhuri & Fiete, 2016). In these models when a to-be-remembered stimulus is presented, it activates a specific population of neurons that remain firing at an elevated rate for as long as necessary until the information is no longer required (Constantinidis et al., 2018). A great deal of work in computational neuroscience has developed mechanisms for sustained stimulus-specific firing at the level of circuits, channels, and LFPs (Compte et al., 2000; Durstewitz et al., 2010; Chaudhuri & Fiete, 2016).

However, there are both theoretical concerns that raise challenges to simple persistent activity models of memory. Modeling work shows that persistent activity is metabolically expensive (Lundqvist, Herman, & Miller, 2018; Stokes, 2015). In attractor models of persistent firing, even a small, temporary disruption to the network can disrupt the memory (Lundqvist et al., 2018). Of particular relevance to temporal continuity (discussed in the next section), information about the passage of time is lost because the firing rate is constant while the stimulus is maintained in working memory. Thus a memory representation based on sustained firing is not sufficient to represent information about time or maintain temporal continuity. These modeling and theoretical objections do not rule out the possibility of some role of persistent firing in working memory, but they do indicate that other forms of memory encoding are needed.

Empirical results have directly shown activity beyond simple persistent firing in memory. Several studies have proposed and identified evidence for various dynamic coding schemes. Stokes (2015) proposed “activity-silent” dynamic coding in which the memory does not depend on continuous neural activity but rather modulation of synaptic weights. Activity-silent working memory would address theoretical concerns about the metabolic cost of working memory. Sreenivasan, Curtis, and D’Esposito (2014) found evidence to suggest that lPFC supports more goal directed information and uses dynamic coding in addition to persistent activity. Sreenivasan and D’Esposito (2019) reviewed numerous examples of dynamic coding, and found evidence that LFP bursts play a role in working memory. Collectively these results were not mutually exclusive with persistent activity serving a role in memory, but they do indicate that memory in the brain involves a wider variety of neural activity. Our results are consistent with some conceptions of dynamic coding. Collectively, time cell activity is equivalent to a smooth trajectory through a state space.

Several studies have presented explanations for why persistent firing has been identified over other more complex actitvity patterns. Lundqvist et al. (2018) argued that more complex tasks result in more complex neural activity, and that previous studies have used simpler tasks, resulting in observations of simple persistent firing. Shafi et al. (2007) proposed that persistent activity could be a result of trial averaging that could be obscuring more complex neural activity and showed substantial variability in firing frequency across trials. Sreenivasan et al. (2014) suggested that sensory cortex maintains the representation while other areas support other forms of coding such as goal directed information. In this study, time cells with sufficiently high constant terms or wide time fields could be potentially misidentified as persistent firing cells by a less thorough analysis.

The time cells reported here were consistent with models of working memory that propose continuous delay duration activity without stable persistent activity by individual neurons. In particular, these time cells were most consistent with models of working memory that have proposed sequentially active firing (Goldman, 2009; Grossberg & Merrill, 1992). The existence of time cells does not directly address the predictions of activity-silent models of working memory, but is partially consistent with these models. In particular, each time cell is active at particular times within a duration, and may be silent during the rest of the duration. The existence of time cells does not address the idea that working memory is stored in functional connectivity, one of the primary prediction of activity-silent models (Stokes, 2015). The population level analysis directly contradicted strictly persistent firing, because persistent firing alone would result in a decoding accuracy that is constant across the entire interval instead of higher at the diagonals. These results do not rule out the possibility that other populations of neurons maintain persistent activity, or that this population of neurons might maintain persistent activity in response to other tasks, but they do show there is a population of neurons in HPC and PFC that do not need persistent activity to encode stimulus identity. The findings in this paper are very much in line with the predictions of a model of working memory maintenance in which the brain estimates the temporal history of past events as a compressed timeline (Howard, Shankar, Aue, & Criss, 2015; Singh, Tiganj, & Howard, in press; Howard, 2018).

### Temporal Continuity in Vision

Temporal continuity has been proposed as a key mechanism in the construction of invariant object representations (DiCarlo, Zoccolan, & Rust, 2012). Invariant object representation refers to the hypothesis that in visual object recognition the brain is able to recognize objects at a variety of viewing angles, distances, and other conditions by constructing a representation that is constant across all these conditions. Because objects are usually temporally continuous with themselves, even as the viewing conditions change, temporal continuity could be used as a cue in learning invariant representations. For example, walking towards an object, you sequentially see it appearing larger and larger as you get closer and closer. In order for the brain to utilize temporal continuity, it needs a temporal signal with object identity information that spans the relevant time scales. Stimulus specific time cells with the collective proprieties of a compressed representation are a suitable case of such a temporal signal. Temporal continuity might play a role not only in building invariant representations, but also in statistical learning and other forms of associative learning and memory across multiple timescales. This is suggested by empirical results on timescales of saccades (Li & DiCarlo, 2008), seconds (Schapiro, Rogers, Cordova, Turk-Browne, & Botvinick, 2013), and tens of seconds (Miyashita, 1988). The particular role of HPC and PFC in visual associative memory is suggested by the underlying neuroanatomy, including lesioning studies (Higuchi & Miyashita, 1996; Schapiro, Turk-Browne, Norman, & Botvinick, 2016) and human fMRI studies (Schapiro, Kustner, & Turk-Browne, 2012; Jackson III & Schacter, 2004).

Empirical results show temporal continuity causes neural representations to become similar for visual stimuli across a variety of scales. In Li and DiCarlo (2008) monkeys were presented with stimuli that were consistently swapped during saccade. This procedure, with repeated exposures, caused neurons in the inferior temporal cortex (ITC) selective for one stimulus over another to become equally selective to both stimuli being exchanged. The stimulus before switching is temporally continuous to the stimulus after switching, and their neural representations become similar. In Schapiro et al. (2013) stimuli are presented continually to human subjects, with their order dictated by a random walk through a community structure that was not known to the participants. fMRI activation patterns for stimuli adjacent in the structure become more similar. In other words, a neural representation become similar according to temporal continuity. In Miyashita (1988) sequentially presented stimuli develop correlations in their neural representations in the ITC through successive presentation. In all of these results, the continuity between visual stimuli presentation causes neural representations (whether neural firing in the case of (Li & DiCarlo, 2008; Miyashita, 1988) or fMRI activation patterns in the case of (Schapiro et al., 2013)) to become more similar.

The smoothly-changing temporal signal observed in HPC and PFC in this study has many of the properties required to enable temporal continuity to influence visual representations. Indeed, there is a wealth of data that implicates HPC and PFC in vision. HPC and PFC are both connected to areas responsible for visual processing. The HPC is reciprocally connected to the EC which is in turn connected to the inferior temporal cortex (ITC) (Canto, Wouterlood, & Witter, 2008). The PFC (area 46 in particular) is reciprocally connected to the entorhinal cortex (EC) and perirhinal cortex (PRC) (Yeterian, Pandya, Tomaiuolo, & Petrides, 2012). The PFC (areas 45 and 46) is also reciprocally connected to the ITC (area TE in particular) (Kravitz, Saleem, Baker, Ungerleider, & Mishkin, 2013). The ITC is one of the last stages of the visual ventral stream, the network of brain regions responsible for object identification. Lesioning monkey EC and PFC eliminated visual associative coding in inferior temporal cortex (ITC), while leaving the overall response to visual stimuli intact (Higuchi & Miyashita, 1996). (Naya, Yoshida, & Miyashita, 2001) showed that the associative signal originates past the ventral stream, possibly in the HPC and PFC, and propagates backwards to the ITC. Miller, Erickson, and Desimone (1996) showed visual stimulus selective neurons exist in the PFC in associative learning. Miller et al. (1996) also showed that compared with the ITC, these PFC neurons showed a stronger response to learned associations, indicating that visual associative learning is present in PFC. Collectively, this evidence suggests that the underlying connectivity could allow the temporal signal we identified in HPC and PFC to propagate to the ITC and contribute to forming associative relationships in the visual system.

Indeed, many studies implicate HPC and PFC in visual statistical learning. In a case study of a patient with complete bilateral hippocampal loss and broader medial temporal lobe (MTL, of which the HPC, EC, and PRC are subregions) damage, the patient was able to discriminate which individual visual shape and scene stimuli they had been exposed to, but was unable to distinguish novel sequences from recurrent sequences of the visual stimuli, indicating an inability to perform statistical learning (Schapiro et al., 2016). Human fMRI studies show the role of MTL (Schapiro et al., 2012; Jackson III & Schacter, 2004) and PFC (Ranganath, Cohen, Dam, & D’Esposito, 2004) in associate learning and statistical learning. Collectively, all of these results suggest that HPC and PFC have an important role in visual statistical learning.

Overall, the evidence suggests that a temporal signal sufficient to support temporal continuity exists in HPC and PFC, that temporal continuity supports associative memory (Schapiro et al., 2013; Miyashita, 1988), and that HPC and PFC are important to associative memory (Higuchi & Miyashita, 1996; Schapiro et al., 2016, 2012; Jackson III & Schacter, 2004). The proprieties of time cells reported here are consistent with the proprieties necessary for constructing associative representations across a variety of scales.

