## Supplemental and Supporting Information for "Conjunctive representation of what and when in monkey hippocampus and lateral prefrontal cortex during an associative memory task"

Nathanael A. Cruzado, Zoran Tiganj
Boston University
Scott L. Brincat, Earl K. Miller
Massachusetts Institute of Technology
Marc W. Howard
Boston University

#### Overview

This supplemental includes additional data, figures, and exploration of the methods of the main text including analysis with a broader criteria for time cells (including cells classified as ambiguous cells in the main text), an alternate method for identifying monotonically changing cells, qualitative review of plots of monotonically changing cells, and analysis and figures of ideal simulated cells. Repeating analyses from the main text on the broader set of time cells gives a similar overall conclusion of temporal compression. An alternative method for identifying monotonically changing cells (using a linear function of firing rate) did not generate substantially different results than those of the main text which used a broad range of Gaussian parameters to approximate linear firing. Qualitative analysis of the raster plots for monotonically changing cells suggests they were not as reliable or systemic as time cells. Simulated time cells with varying combinations of ideal properties show how the analysis methods handle data with and without various forms of temporal compression.

#### Ambiguous Cells

For the purpose of getting a conservative count of time cells, a restrictive criteria for time cells was used in the main text. However, the strictness of this criteria might have also biased the collective characteristics of these time cells. Thus the analyses of the main text are repeated with a broader criteria in this supplemental. Broadening the criteria for time cells does reduce the magnitude of their temporal compression, but temporal compression is still present.

##### Definition and Comparison

Cells for which the Gaussian receptive field fit better than a constant firing rate and peaked within the 0-1.25 second time window, but were within one standard deviation of either the 0 second lower limit or 1.25 second upper limit were classified as ambiguous cells. Ambiguous cells were excluded from the primary analyses and figures of the main text. For this supplemental, the analyses techniques and plots of Figure 4 are repeated with ambiguous cells in Figure S1. Overall, compared to Figure 4, in Figure S1 the heatmaps look almost the same, the temporal compression in peak time versus width is reduced but still present, and the temporal compression is present at the same statistical significance in the CDFs.

#### Results

Figure S1 has two features demonstrating temporal compression. First, time fields later in the delay were more broad than time fields earlier in the delay. This can be seen as the widening of the central ridge as the peak moves to the right. This is also shown in plots of the relationship between time field peak time and width (Figure S1c and S1e). For HPC this relationship is positive, but only weakly significant (intercept $.15 \pm .026$ s $p < .001$, slope $.07 \pm .039$ s/s $p < .1$). For PFC this relationship is positive and significant (intercept $.06 \pm .017$ s $p < .001$, slope $.06 \pm .027$ s/s $p < .05$, $r^2 = .16$). Second, peak times of the time cells were not evenly distributed across the delay, with later time periods represented by fewer cells than early time periods. This can be seen in the curvature of the central ridge; a uniform distribution of time fields would manifest as a straight line. This is also shown in peak time CDF (Figure S1d and S1f). For both regions, the peak time distributions differs from a uniform distribution (HPC: $n = 159$, ks-stat=0.18, $p < .001$; PFC: $n = 320$, ks-stat=0.09, $p < .001$). Overall, although including ambiguous cells reduces the magnitude of temporal compression, temporal compression is still present.

### Monotonically Changing Cells

To further explore monotonically changing cells, an alternative method for identifying them was implemented and a simulated data set of ideal time cells with varying properties was analyzed. Including monotonically changing cells identified by this alternate method in the overall count of monotonically changing cells does not increase their number enough to alter the overall conclusion that time cells were more common than monotonically changing cells.

#### Methods for Alternative Analysis

In order to further model ramping and decaying firing rates, two additional linear function were optimized to fit firing rates using the maximum likelihood parameter estimation described in the main text. These additional linear function proved largely redundant with the Gaussian receptive field used in the main text. One of these linear functions is intended to fit cells with a monotonically increasing firing rate, the other is intended to fit cells with a decaying firing rate. As in the methods of the main text, each of these functions represents the probability of firing within each millisecond time bin.

$$p\,(t;\Theta) = a_0 + a_1 t/1000 \tag{S1}$$

$$p\,(t;\Theta) = a_0 + a_1(1.25 - t/1000) \tag{S2}$$

Equation (S1) describes a ramping firing rate, in which as each additional time bin elapses, the probability of firing increases by $a_1/1000$. Equation (S2) describes a decaying

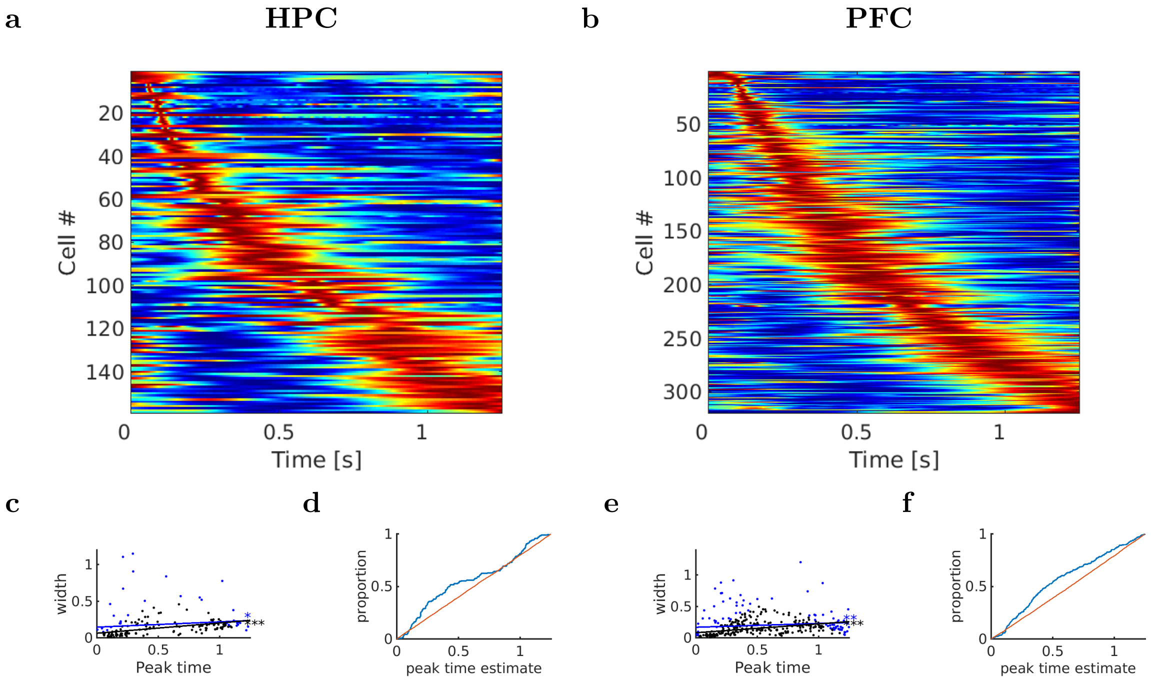


*Supplementary Figure S1*. **Using a less stringent threshold for including cells in the analysis, the figure and results come out nearly identical to Figure 4** Activity of 159 HPC units and 320 PFC units classified as time cells or ambiguous cells. Each row corresponds to a single unit and displays the firing rate (normalized to 1) averaged across all trials. Red corresponds to high firing rate; blue corresponds to low firing rate. The time cells are sorted with respect to the peak time estimated for their time field. There are two features related to temporal accuracy that can be seen from examination of these figures. First, time fields later in the delay were more broad than time fields earlier in the delay (see also **c** and **e**). This can be seen as the widening of the central ridge as the peak moves to the right. In addition the peak times of the time cells were not evenly distributed across the delay, with later time periods represented by fewer cells than early time periods (see also **d** and **f**). This can be seen in the curvature of the central ridge; a uniform distribution of time fields would manifest as a straight line. **c:** Width of the time fields as a function of the peak time in HPC. Each dot represents the best-fitting parameters for a single unit classified as a time cell. The blue line is a fitted linear model for both time and ambiguous cells. The black line is a fitted linear model for only time cells. The linear regression is significant ($p < .001$) when only using time cells, and is only weakly significant, $p < .1$ when including ambiguous cells (blue dots). **d:** Peak times of the time fields in HPC are non-uniformly distributed along the delay interval, even when including ambiguous cells. The blue line is the cumulative distribution function of the time cell peak times. The red line is a uniform distribution. The distributions differ at $p < .001$ as estimated from a Kolmogorov-Smirnov test. **e:** Same as **c** but for time cells and ambiguous cells in PFC. The linear regression is significant ($p < .001$), and less significant when including ambiguous cells ($p < .05$). **f:** Same as **d** but for time cells in PFC. Kolmogorov-Smirnov test is significant at $p < .001$. Overall, adding the ambiguous cell into the analysis did not affect the overall results.

firing rate, in which as each additional time bin elapses, the probability of firing decreases by $a_1/1000$. These equations can also be conceptualized in terms of average firing rate. For Equation (S1), initial firing rate is at $1000a_0$ Hz and as each additional millisecond elapses firing rate increases by $a_1$ Hz. For Equation (S2), initial firing rate is at $1000(a_0+a_1 1.25)$ Hz and as each additional millisecond elapses, firing rate decreases by $a_1$ Hz. In order to ensure that $p\,(t;\Theta)$ can be considered as a probability we had to ensure that its values are bounded between 0 and 1. Therefore, the coefficients were bounded such that $a_0+a_1 1.25 \leq 1$. Neither $a_0$ or $a_1$ were allowed to go below 0.

These linear firing rates are equivalent to the constant rate when $a_1$ is equal to zero. Therefore the linear models are nested with the constant model and can be compared with a likelihood-ratio test. However, the linear firing rates are not nested with the Gaussian receptive field. Therefore a different approach is needed to directly compare them. The Akaike Information Criterion (AIC) was used to compare fits of the linear model to the Gaussian receptive field model. Cells fit better by one of these linear functions than a constant rate firing rate or a Gaussian receptive field were classified as monotonically changing cells.

**Results.** No time cells with $\mu$ within the 0-1.25 second bound were better fit by a linear firing rate than a Gaussian receptive field. Thus, this additional analysis did not change the results regarding time cells or ambiguous cells. These linear models only introduced a small additional number of cells to be classified as monotonically changing cells (13 more cells for HPC, for 27 total and 19 more cells for PFC for 41 total). Even with these additional cells, the proportion of monotonically changing cells is still significantly less than the proportion of time cells (significant at p<.01 by a two proportion z-test).

**Why additional modeling of monotonically changing cells proved redundant.** Given the large range of values the Gaussian receptive field was allowed to take, the Gaussian receptive field was able to accurately approximate linearly changing firing rates (note how the models exactly overlap in Figure S2c, S3a, and S3b and are close for the other example cells). This is further illustrated in Figure S4.

#### Example Cells

Some of the best examples of monotonically changing cells are shown in Figure S2 and Figure S3. For each example cell rasters, averaged firing rates, fitted Gaussian receptive fields, and fitted linear functions are shown. These plots show various irregularities that illustrate that monotonically changing cells were not as reliable as time cells. Even among these best examples there are irregularities. One of these examples cells (Figure S2d) was not firing across all trials, indicating possible recording or spike sorting issues. Two example cells (Figure S2b and S3d) showed changes in firing rate across trials. Four example cells (Figure S2a, S2c, S3b, and S3d) showed deviations in average firing rate which might suggest other forms of temporal modulation in addition or in place of monotonically changing firing rates. For instance, periodic oscillation over the delay interval might be misclassified as a monotonically changing firing rate. These oscillations might also simply be noise. Although these were the best examples of monotonically changing cells, they show irregularities, indicating that the monotonically changing cells are not as reliable or consistent as the time cells.

Overall, there was significantly less monotonically changing cells than time cells, even including monotonically changing cells identified with an additional methodology and including monotonically changing cells that show irregularities in their rasters. Thus, time cells were the primary form of temporal coding in PFC and HPC.

#### Ideal Simulated Data

In order to further explore the performance of the maximum likelihood time cell estimation and of the analysis techniques meant to address temporal compression, four populations of ideal time cells were simulated: temporally compressed firing field width and temporally compressed peak times, uniform firing field width and temporally compressed peak times, temporally compressed firing field widths and uniformly spread peak times, uniform firing field width and uniformly spread peak times. Analyses of simulated time cell peak time versus firing field width and peak time CDFs generated results consistent with the expected results based on the simulated data properties.

**Simulation Methods.** Each population was simulated by first analytically calculating a firing rate for each simulated cell according to the desired properties. Then, for each simulated millisecond, the probability of firing was set using this normalized simulated firing rate, with a maximum firing rate selected for plausibility with empirical spiking data (peaking at 10 Hz).

The firing rates for simulated time cells with temporally compressed firing field width and temporally compressed peak times were calculated using Equation (S3). The values of $\overset{*}{\tau}_i$ were geometrically spaced (with each successive $\overset{*}{\tau}_i$ increased by a constant ratio) with peaks from .02 second to 1.5 seconds.

$$\tilde{f}(t) \propto \frac{1}{\overset{*}{\tau}_i} \left( \frac{t}{\overset{*}{\tau}_i} \right)^k e^{-\frac{kt}{\overset{*}{\tau}_i}} \tag{S3}$$

The firing rates for simulated time cells with uniform firing field width and temporally compressed peak times were calculated as Gaussian curves with standard deviation equal to 200 ms, and values of $\overset{*}{\tau}_i$ were geometrically spaced (with each successive $\overset{*}{\tau}_i$ increased by a constant ratio) with peaks from .02 second to 1.5 seconds (Equation (S4)).

$$\tilde{f}(t) \propto e^{\frac{-(t-\overset{*}{\tau}_i)^2}{2\sigma_t^2}} \tag{S4}$$

The firing rate for simulated time cells with temporally compressed firing field widths (proportional to half their peak time) and uniformly spread peak times (from .015 seconds to 1.5 second in .015 second steps) were calculated using Equation (S4).

The firing rate for simulated time cells with uniform firing field width (200 ms standard deviation) and uniformly spread peak times (uniformly spaced from .015 seconds to 1.5 seconds in .015 second steps) were calculated using Equation (S4).

##### Simulation Results

**Compressed Firing Field Width, Compressed Peak Time.** For simulated time cells with temporally compressed firing field width and temporally compressed peak

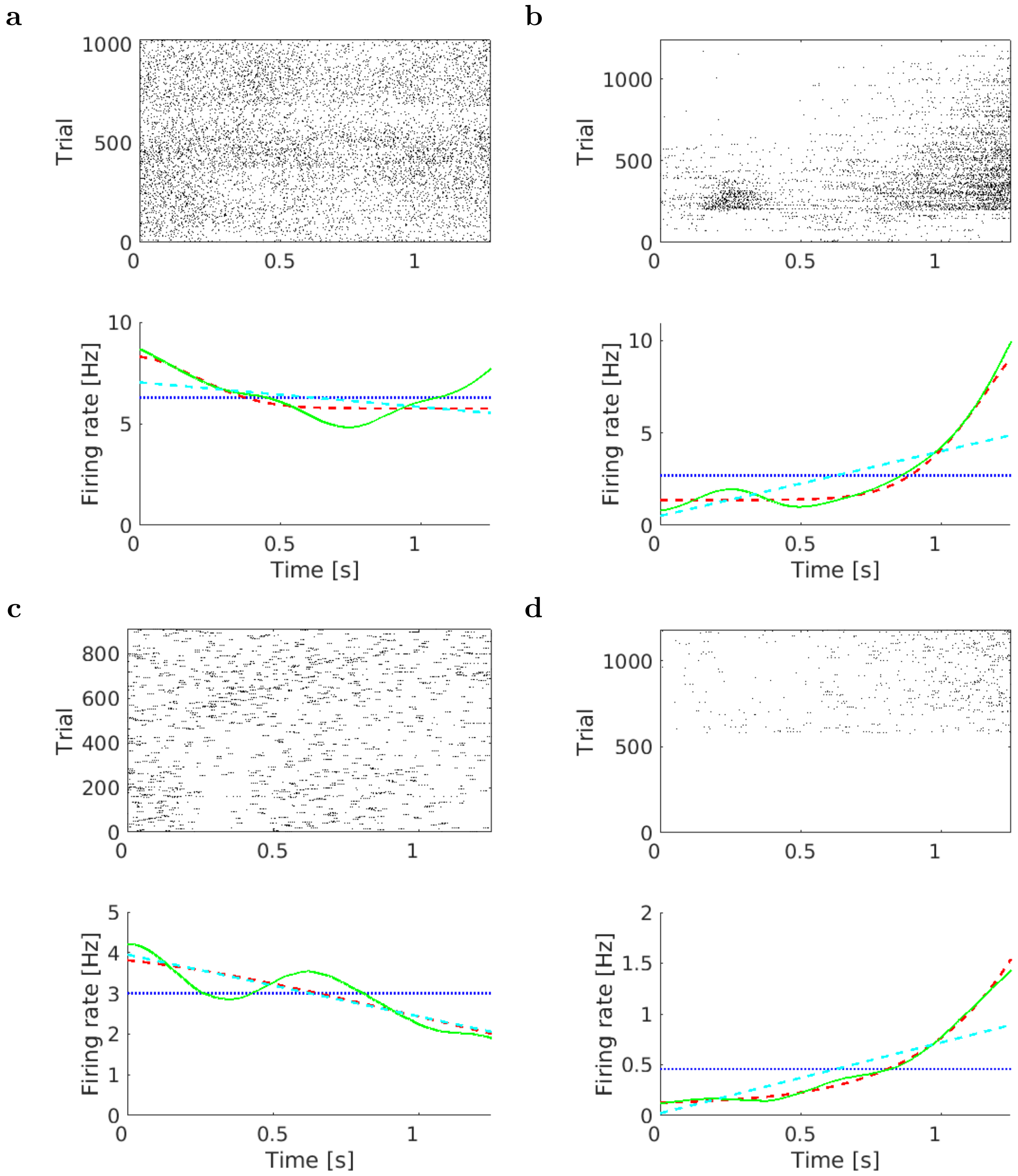


*Supplementary Figure S2*. **Examples of HPC cells that exhibit possible monotonically changing firing rate** Within each raster each row is a trial, and time is shown from cue stimulus onset to associate stimulus onset. Within each graph, the green line is the smoothed firing rate, the blue line is a fitted constant firing rate, the red line is the fitted Gaussian, and the blue line is a fitted linear firing rate. Note that for **c**, the linear firing rate and Gaussian firing rate overlap exactly, and that the two models also give close prediction for the other cells. In **a** the cell was classified as monotonically changing because the firing rate decreased overall, however actually looking at the smoothed firing rate, it might possible be doing something more complicated, as it seem to increase after .75 seconds. **b**, the firing rate is ramping up, however this might be an artifact of the cell changing its firing field width across trials (starting with a wide firing field in the first 500 trials and then narrowing in the later 500 trials). **c** shows a decaying firing rate. **d** shows an increasing firing rate, however the cell seems to cut in at around 500 trials, possibly as a spike sorting artifact or as an artifact of the recording electrode shifting. Qualitative inspection of possible monotonically changing cells beyond the 0-1.25 second range reveals a variety of possible forms of temporal activity: time cell like peaking, continued ramping, and sudden changes in firing rate similar. Overall these plots cannot be treated as good evidence of monotonically changing cells.

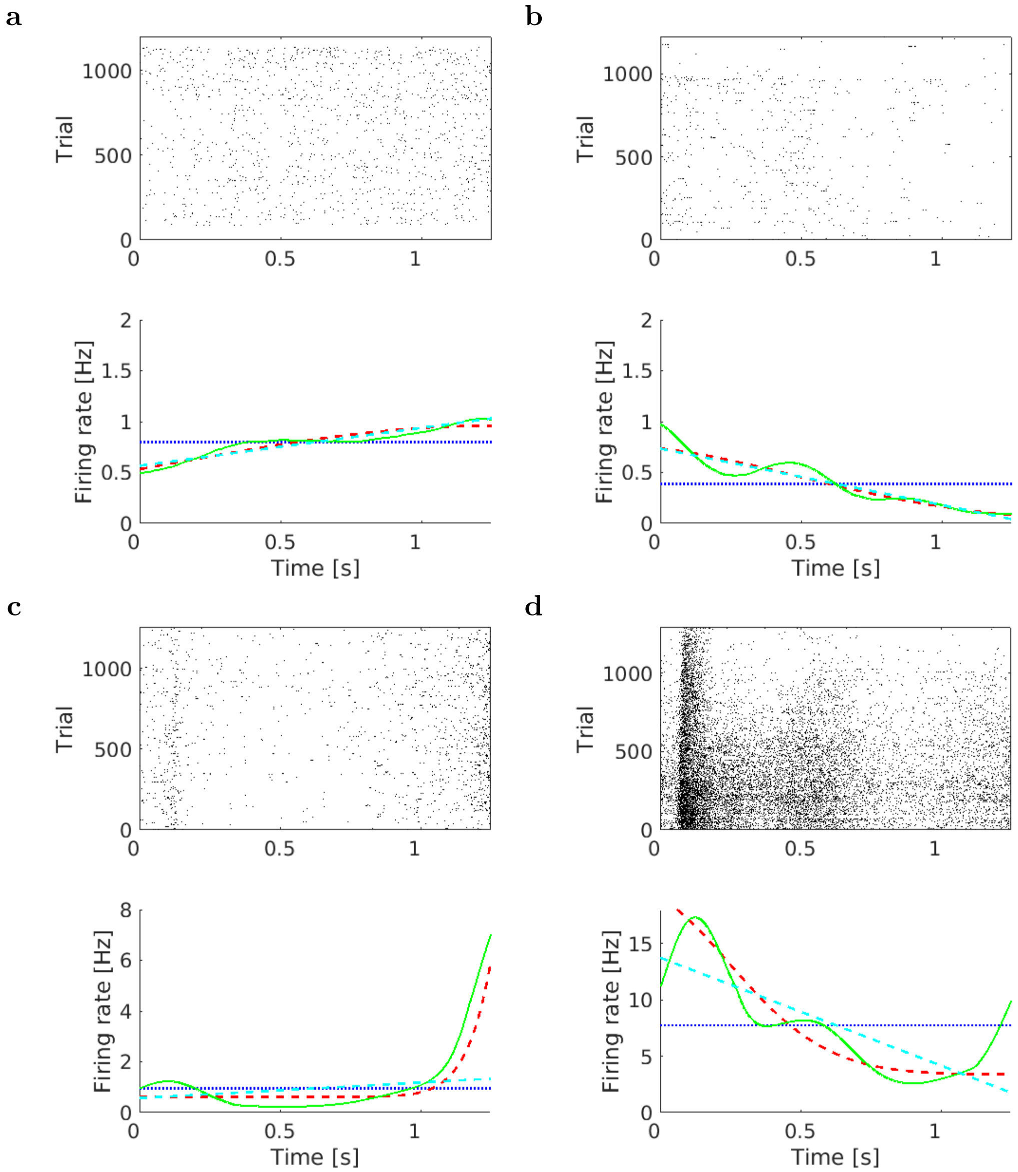


*Supplementary Figure S3*. **Examples of PFC cells that met criteria for monotonically changing firing rate** Within each raster each row is a trial, and time is shown from cue stimulus onset to associate stimulus onset. Within each graph, the green line is the smoothed firing rate, the blue line is a fitted constant firing rate, the red line is the fitted Gaussian, and the blue line is a fitted linear firing rate. Note that for **a** and **b**, the linear firing rate and Gaussian firing rate overlap exactly, and that the two models also give close prediction for the other cells. In **a** firing rate is ramping up. **b** firing rate is decaying. **c** firing rate ramps up suddenly at the end of the 1.25 second interval. **d** has a rapid increase in firing rate at around .1 seconds and then drops off. This could possibly be closer to a time cell with skewed firing field than a decaying firing rate. Conceptually speaking, much of the literature doesn't precisely distinguish between these concepts. Qualitative inspection of possible monotonically changing cells beyond the 0-1.25 second range reveals a variety of possible forms of temporal activity: time cell like peaking, continued ramping, and sudden changes in firing rate similar. Overall these plots cannot be treated as good evidence of monotonically changing cells.

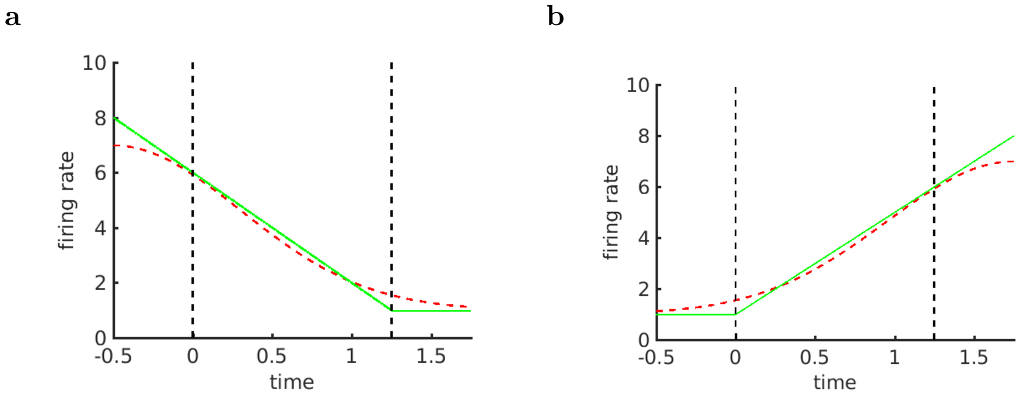


*Supplementary Figure S4*. **Ramping and decaying firing rates can be described by the time cell model**. With the appropriate parameters, linear functions of firing rate (green lines) can be approximated with a Gaussian receptive field(dotted red lines) if its parameters are allowed to take the appropriate values. Note that the likelihood of the fit is only calculated for the 0 to 1.25 second range, identified with dotted black lines.

times, 87 cells were identified as time cells and 11 cells were identified as ambiguous cells. The linear regression for peak time vs. width gave results: using only time cells: intercept $-.021 \pm .005$ $p < .001$, slope $.16 \pm .01$ $p < .001$, $r^2 = .87$; including ambiguous cells and time cells: intercept $-.05 \pm .01$ $p < .001$, slope $.23 \pm .01$ $p < .001$, $r^2 = .83$). The KS test rejected the hypothesis that the distribution of the peak times is uniform (using only time cells $n = 87$, ks-stat=0.44, $p < .001$; using time cells and ambiguous cells $n = 98$, ks-stat=0.35, $p < .001$). Thus these analyses correctly identified the temporally compressed firing field and temporally compressed peak times.

**Uniform Firing Field Width, Compressed Peak Time.** For simulated time cells with uniform firing field width and temporally compressed peak times, 40 cells were identified as time cells and 55 cells were identified as ambiguous cells (a large number of cells early in the interval were identified as ambiguous because their firing field width was too close to the bound). The linear regression for peak time vs. width gave results: using only time: cells intercept $.20 \pm .002$ $p < .001$, slope $.00 \pm .003$ $p = .81$, $r^2 = .002$; including ambiguous cells and time cells: intercept $.20 \pm .0001$ $p < .001$, slope $-.003 \pm .002$ $p < .1$, $r^2 = .033$). The KS test rejected the hypothesis that the distribution of the peak times is uniform (using only time cells $n = 40$, ks-stat=0.20, $p < .1$; using time cells and ambiguous cells $n = 95$, ks-stat=0.38, $p < .001$). Thus these analyses correctly identified the uniform firing field width and temporally compressed peak times.

**Compressed Firing Field Width, Uniform Peak Time.** For simulated time cells with temporally compressed firing field widths and uniformly spread peak times, 70 cells were identified as time cells and 14 cells were identified as ambiguous cells. The linear regression for peak time vs. width gave results: using only time cells: intercept $.000 \pm .0001$ $p < .1$, slope $.49 \pm .002$ $p < .001$, $r^2 = .998$; including ambiguous cells and time cells: intercept $.0023 \pm .0011$ $p < .05$, slope $.49 \pm .003$ $p < .001$, $r^2 = .998$). The KS test rejected the hypothesis that the distribution of the peak times is uniform when using only time cells $n = 70$, ks-stat=0.21, $p < .001$ (the ambiguous cells not included altered the distribution enough to make it seem non-uniform to the analysis). However using time cells and ambiguous cells the KS-test failed to reject the hypothesis that the distribution was uniform $n = 84$, ks-stat=0.06, $p = .94$). Qualitatively, looking at the CDF (Figure S5), it is clear that the distribution is very close to uniform. Thus these analyses correctly identified the temporally compressed firing field and had some slight issues correctly identifying the uniformly spread peak times.

**Uniform Firing Field Width, Uniform Peak Time.** For simulated time cells with uniform firing field widths and uniformly spread peak times, 55 cells were identified as time cells and 29 cells were identified as ambiguous cells (many of the cells at the beginning and end of the interval were identified as ambiguous). The linear regression for peak time vs. width gave results: using only time cells: intercept $.20 \pm .001$ $p < .001$, slope $-.002 \pm .002$ $p = .18$, $r^2 = .033$; including ambiguous cells and time cells: intercept $.20 \pm .001$ $p < .001$, slope $-.003 \pm .002$ $p < .05$, $r^2 = .049$). The KS test reject the hypothesis that the distribution of the peak times was uniform when using only time cells because cells near the edge are all excluded as ambiguous cells ($n = 55$, ks-stat=0.22, $p < .001$) but including ambiguous cells, the KS test fails to reject the null hypothesis $n = 84$, ks-stat=0.028, $p \approx 1$). Qualitatively, looking at the CDF including time cells and ambiguous cells (Figure S5 m), it is clear that the distribution is uniform. Thus these analyses correctly identified the uniform firing field widths but failed to identify the uniformly spread peak times when looking at only time cells, but an analysis including ambiguous cells and a qualitative examination of the resulting plots makes it clear that the peak time distribution is uniform.

Thus the analysis techniques for identifying temporal compression correctly identified temporal compression (or lack thereof) in both peak times vs. firing field widths and peak time CDFs in 4 sets of simulated data. Notably, including ambiguous cells was necessary for the analyses of time cells with uniform firing field width and uniform peak time. This is because the firing field width was set wide enough that a large number of cells were identified as ambiguous cells in a way that would alter the overall distribution of time cell peak times.

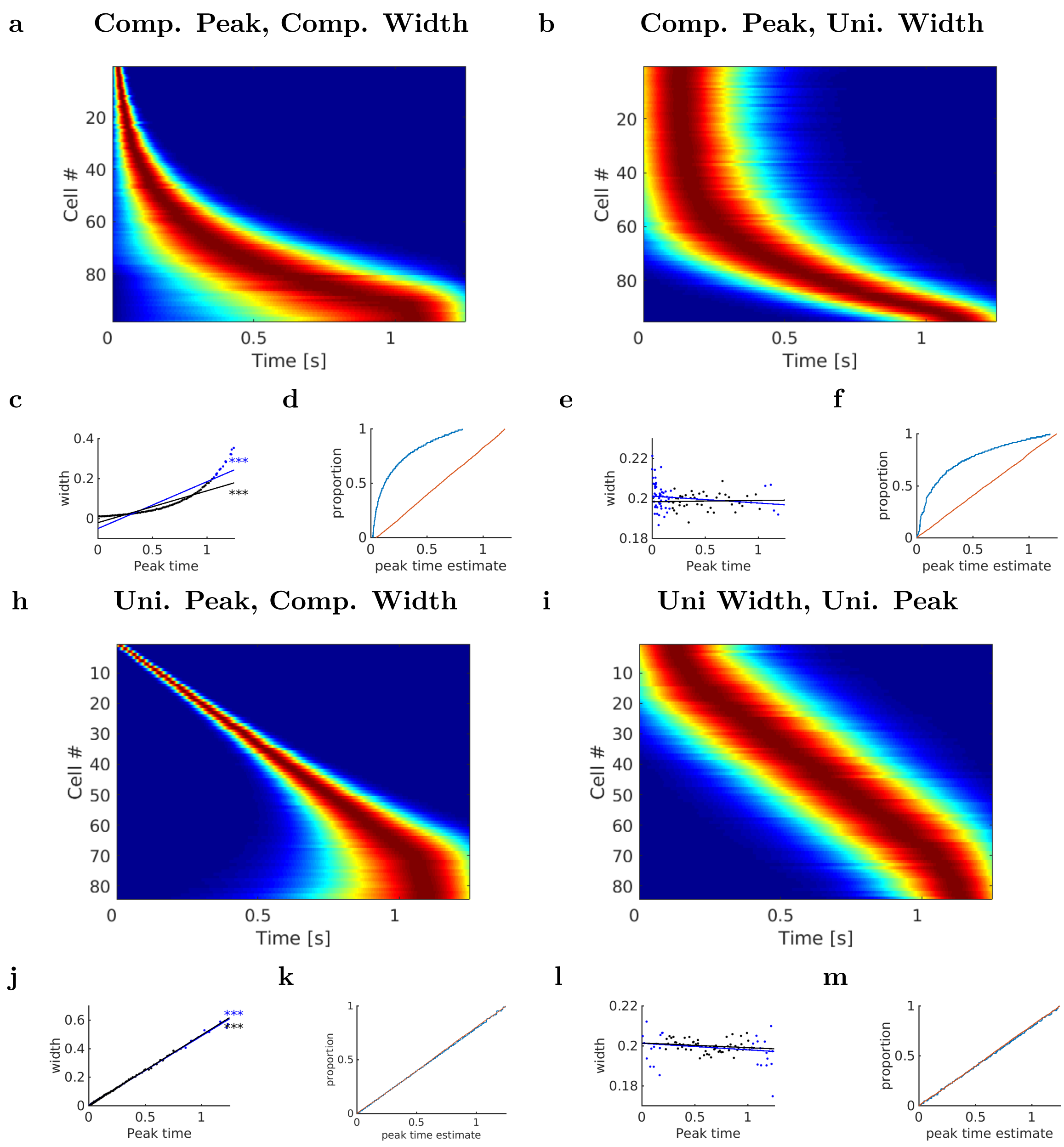


*Supplementary Figure S5*. **Ideal simulated cells with various properties illustrate the performance of the analyses techniques.** Analyses and figure design is as in main text Figure 4 and Supplemental Figure S1. 100 time cells were simulated with compressed peaks or uniform peaks and compressed width or uniform widths. Compressed peaks were spaced out according to a power law such that each peak was a set ratio from the previous peak. Uniform peaks were spaced evenly .0125 seconds apart. Uniform widths were set to 200 ms. All plots include both time cells and ambiguous cells. **a, c, d** Compressed peak time and compressed widths. **b, e, f** compressed peaks and uniform width. **h, j, k** uniform peaks and compressed width. **i, l, m** uniform peaks and uniform widths. Analysis of peak time vs. firing field width **c,e,j,l** and of the CDFs of peak times **d,f,k,m** generated the expected results in identifying compressed vs. uniform properties. Note the heatmaps **a,b,h,i** have slight distortion in appearance in the edges due to Gaussian smoothing.
